# Rapid low-cost assembly of the *Drosophila melanogaster* reference genome using low-coverage, long-read sequencing

**DOI:** 10.1101/267401

**Authors:** Edwin A. Solares, Mahul Chakraborty, Danny E. Miller, Shannon Kalsow, Kate Hall, Anoja G. Perera, J.J. Emerson, R. Scott Hawley

**Author notes:** Department of Pediatrics, Seattle Children’s Hospital and University of Washington, 4800 Sand Point Way NE, Seattle, WA, 98105, USA. These authors contributed equally to this work (Edwin A. Solares, Mahul Chakraborty). 0000-0002-3220-4927 (EAS). 0000-0002-6478-0494 (RSH). 0000-0003-2414-9187 (MC). 0000-0001-9474-0891 (JJE).

## Abstract

Accurate and comprehensive characterization of genetic variation is essential for deciphering the genetic basis of diseases and other phenotypes. A vast amount of genetic variation stems from large-scale sequence changes arising from the duplication, deletion, inversion, and translocation of sequences. In the past 10 years, high-throughput short reads have greatly expanded our ability to assay sequence variation due to single nucleotide polymorphisms. However, a recent *de novo* assembly of a second *Drosophila melanogaster* reference genome has revealed that short read genotyping methods miss hundreds of structural variants, including those affecting phenotypes. While genomes assembled using high-coverage long reads can achieve high levels of contiguity and completeness, concerns about cost, errors, and low yield have limited widespread adoption of such sequencing approaches. Here we resequenced the reference strain of *D. melanogaster* (ISO1) on a single Oxford Nanopore MinION flow cell run for 24 hours. Using only reads longer than 1 kb or with at least 30x coverage, we assembled a highly contiguous *de novo* genome. The addition of inexpensive paired reads and subsequent scaffolding using an optical map technology achieved an assembly with completeness and contiguity comparable to the *D. melanogaster* reference assembly. Comparison of our assembly to the reference assembly of ISO1 uncovered a number of structural variants (SVs), including novel LTR transposable element insertions and duplications affecting genes with developmental, behavioral, and metabolic functions. Collectively, these SVs provide a snapshot of the dynamics of genome evolution. Furthermore, our assembly and comparison to the *D. melanogaster* reference genome demonstrates that high-quality *de novo* assembly of reference genomes and comprehensive variant discovery using such assemblies are now possible by a single lab for under $1,000 (USD).

## INTRODUCTION

The characterization of comprehensive genetic variation is crucial for the discovery of mutations affecting phenotypes. In the last 10 years, the exponential decline in cost and the explosion of high-throughput for short-read sequencing has revolutionized our ability to assay genome-wide sequence variation. Short reads excel at identifying single nucleotide polymorphisms (SNPs) and short insertion-deletion (indel) polymorphisms in unique genomic regions. However, the majority of the sequence difference between individuals is caused by duplications, deletions, inversions, or translocation of sequences—collectively known as structural variants (SVs) (Feuk, Carson, and Scherer 2006). SVs are often longer than short sequencing reads (generally 50-150 bp), meaning genotyping of SVs is indirect and relies on features of alignments to a reference genome such as divergent read mappings, split reads, or elevated read coverage (Medvedev, Stanciu, and Brudno 2009; Alkan, Coe, and Eichler 2011). However, comparisons of extremely contiguous *de novo* genome assemblies from humans (Huddleston and Eichler 2016; M. J. P. Chaisson et al. 2017) and *Drosophila melanogaster* (Chakraborty et al. 2018) revealed that short reads miss 40-80% of SVs. Consequently, methods that are not susceptible to the shortcomings of short-read sequencing are essential to obtain a more complete view of genome variation (Alkan, Coe, and Eichler 2011). We propose one approach—comparison of contiguous and accurate *de novo* genome assemblies—that would overcome these limitations and drastically improve our understanding of genetic variation (Alkan, Coe, and Eichler 2011).

Although short reads have been used extensively for *de novo* genome assembly, they fail to resolve repetitive regions in genomes, leaving errors and gaps while assembling such regions (Paszkiewicz and Studholme 2010; Treangen and Salzberg 2011; Bradnam et al. 2013). Such fragmented draft-quality assemblies are therefore poorly suited for identification of SVs (Alkan, Coe, and Eichler 2011) and lead to incomplete and/or missing gene models (Gordon et al. 2016). Theoretical considerations of the genome assembly problem predict that, with sufficient read depth and length, genome assemblies can resolve even difficult regions (Motahari, Bresler, and Tse 2013; Lam, Khalak, and Tse 2014; Bresler, Bresler, and Tse 2013; Shomorony, Courtade, and Tse 2016). Consistent with this, long reads produced by Single Molecule Real-Time sequencing from Pacific Biosciences (PacBio) and Nanopore sequencing from Oxford Nanopore Technologies (ONT) provide data capable of achieving remarkably contiguous *de novo* genome assemblies (Kim et al. 2014; Berlin et al. 2015; Michael et al. 2018; Jain et al. 2018). However, due to high error rates (∼10-15%), generation of reliable assemblies with these reads requires non-trivial coverage (generally 30x or greater) (Koren et al. 2017; Chakraborty et al. 2016). Nevertheless, until recently, long-read methods have required prohibitively expensive reagents, and technologies like PacBio also require substantial capital investment related to the housing and maintenance of equipment necessary to perform the sequencing. Combined with concerns about high error rates, widespread adoption of long-molecule sequencing for *de novo* assembly and variant detection has been tentative.

Sequencing using ONT may produce reads that are hundreds of kilobases in length (Jain et al. 2018), though their application to *de novo* assembly of reference-grade multicellular eukaryotic genomes is not yet routine. To understand how effective assembly using ONT is when applied to *de novo* genome assembly of a metazoan like *Drosophila,* we measured the contiguity, completeness, and accuracy of a *de novo* assembly constructed with ONT reads. To accomplish this, we resequenced the *D. melanogaster* reference genome strain (ISO1) using the ONT MinION and compared the resulting assembly with the latest release of the *D. melanogaster* reference assembly (Hoskins et al. 2015), which is arguably the best metazoan reference genome available. We followed an assembly merging approach (Chakraborty et al. 2016) to combine modest long-read coverage from a single ONT MinION flow cell (30x depth of coverage with an average read length of 7,122 bp) and Illumina short-read data. This assembly resulted in a highly contiguous and accurate genome assembly. Notably, with this approach, the majority of the euchromatin of each chromosome arm is represented by a single contiguous sequence (contig). Collectively, the assembly recovered 97.7% of Benchmarking Universal Single Copy Orthologs (BUSCOs). This is similar to the 98.3% BUSCOs recovered in the most recent release of the *D. melanogaster* genome (version 6.16). Scaffolding of the assembly with Bionano optical maps led to further improvements in contiguity. Finally, we examined the structural differences between our assembly and the published assembly of the same strain and observed several candidate SVs, the majority of which are transposable element (TE) insertions and copy-number variants (CNVs). These are mutations that must be either: 1) recent mutations that occurred in the genome strain since it was sequenced; 2) segregating in the genome strain due to incomplete isogeny; or 3) errors in one of the assemblies.

Overall, we show that high-quality *de novo* genome assembly of *D. melanogaster* genomes is feasible using low-cost ONT technology, enabling an assembly strategy that can be applied broadly to metazoan genomes. This strategy will make high-quality reference assemblies obtainable for species lacking reference genomes. Moreover, *de novo* assemblies for population samples of metazoan species is now feasible, opening the door for studying evolutionary and functional consequences of structural genetic variation in large populations.

## METHODS

### Stocks

The ISO1 *D. melanogaster* reference stock used for both Nanopore and Illumina sequencing was obtained from the BDGP in 2014 (Hoskins et al. 2015). All flies were kept on standard cornmeal-molasses medium and were maintained at 25°C.

### DNA Isolation and Quantification

DNA for Nanopore sequencing was isolated from males and females using the Qiagen Blood & Cell Culture DNA Mini Kit. Briefly, 60-80 flies were placed in two 1.5-mL Eppendorf Lo-Bind tubes and frozen in liquid nitrogen before being homogenized using a pestle in 250 μL of Buffer G2 with RNAse. 750 μL of Buffer G2 and 20 μL of 20 mg/mL proteinase K was then added to each tube and incubated at 50°C for 2 hr. After 2 hr, each tube was spun at 5,000 RPM for 5 min, and the supernatant was removed and placed in a new 1.5-mL Lo-Bind tube and vortexed for 10 sec. The supernatant from both tubes was then transferred onto the column and allowed to flow through via gravity. The column was washed 3x with wash buffer and eluted twice with 1 mL of elution buffer into two 1.5-mL Lo-Bind tubes. 700 μl of isopropanol was added and mixed via inversion before being spun at 14,000 RPM for 15 min and 4°C. The supernatant was removed and the pellet was washed with 70% ethanol, then spun at 14,000 RPM for 10 min at 4°C. The supernatant was removed and 25 μL of ddH_2_O was added to each tube and allowed to sit at room temperature for 1 hr. Both tubes were then combined into one. DNA was quantified on a Nanodrop and Qubit. DNA for Illumina sequencing was isolated from males and females using the Qiagen DNeasy Blood and Tissue Kit according to the manufacturer’s instructions and quantified on a Qubit.

### Library Preparation, Sequencing, and Basecalling

For Nanopore sequencing, 1.5 μg of DNA (5.49 μL of 273 ng/uL DNA) was used to prepare a 1D sequencing library (SQK-LSK108) according to the manufacturer’s instructions, including the FFPE repair step. The 75-μl library was then immediately loaded onto a R9.5 flow cell prepared according to the manufacturer’s instructions and run for approximately 24 hr. Basecalling was completed using ONT Albacore Sequencing Pipeline Software version 2.0.2. Reads collected during the mux phase of sequencing were not included.

For Illumina sequencing, ∼600-bp fragments were generated from 500 ng of DNA using a Covaris S220 sonicator. Fragments of 500-700 bp were selected using a Pippin and libraries were prepared using a KAPA High Throughput Library Preparation kit and Bioo Scientific NEXTflex DNA Barcodes. The library was pooled with others and run as a 150-bp paired-end run on a single flow cell of an Illumina NextSeq 500 in medium-output mode using RTA version 2.4.11. bcl2fastq2 v2.14 was then run in order to demultiplex reads and generate FASTQ files.

### Genome Assemblies

Canu (Koren et al. 2017) release v1.5 was used to assemble the ONT reads. Canu was run with default parameters in grid mode (Sun Grid Engine) using ONT reads >1 kb and a genome size of 130 Mb. To generate the De Bruijn graph contigs for the hybrid assembly, we used Platanus (Kajitani et al. 2014) v1.2.4 with default settings to assemble 67.4x of Illumina paired-end reads obtained from the DPGP (http://www.dpgp.org/dpgp2/DPGP2.html) (Pool et al. 2012). The hybrid assembly was generated with DBG2OLC (Ye et al. 2016) using contigs from the Platanus assembly and the longest 30x ONT reads. DBG2OLC settings (options: k 25 AdaptiveTh 0.01 KmerCovTh 2 MinOverlap 35 RemoveChimera 1) were similar to those used for PacBio hybrid assembly of ISO1 (Chakraborty et al. 2016), except that the k-mer size was increased to 25 and the MinOverlap to 35 to minimize the number of misassemblies. The consensus stage of DBG2OLC was run with PBDAG-Con (Chin et al. 2013) and BLASR (M. J. Chaisson and Tesler 2012). Separately, minimap v0.2-r123 (using a minimizer size window of 5, FLOAT fraction of minimizers of 0, and min matching length of 100) and miniasm v0.2-r123 (using default settings) were also used to assemble only the ONT reads (Li 2016).

The Canu and DBG2OLC assemblies were merged using quickmerge (Chakraborty et al. 2016). First, the two assemblies were merged using the DBG2OLC assembly as the query and the Canu assembly as the reference. Thus, the first quickmerge run (options: hco 5.0 c 1.5 l 2900000 ml 20000) filled gaps in the DBG2OLC assembly using sequences from the Canu assembly, giving preference to the Canu assembly sequences at the homologous sequence junctions. The contigs that are unique to the Canu assembly were incorporated in the final assembly by a second round of quickmerge. In the second quickmerge run (options: hco 5.0 c 1.5 l 2900000 ml 20000), the merged assembly from the previous step was used as the reference assembly, and the Canu assembly was used as the query assembly (Figure 1).

**Figure 1.**
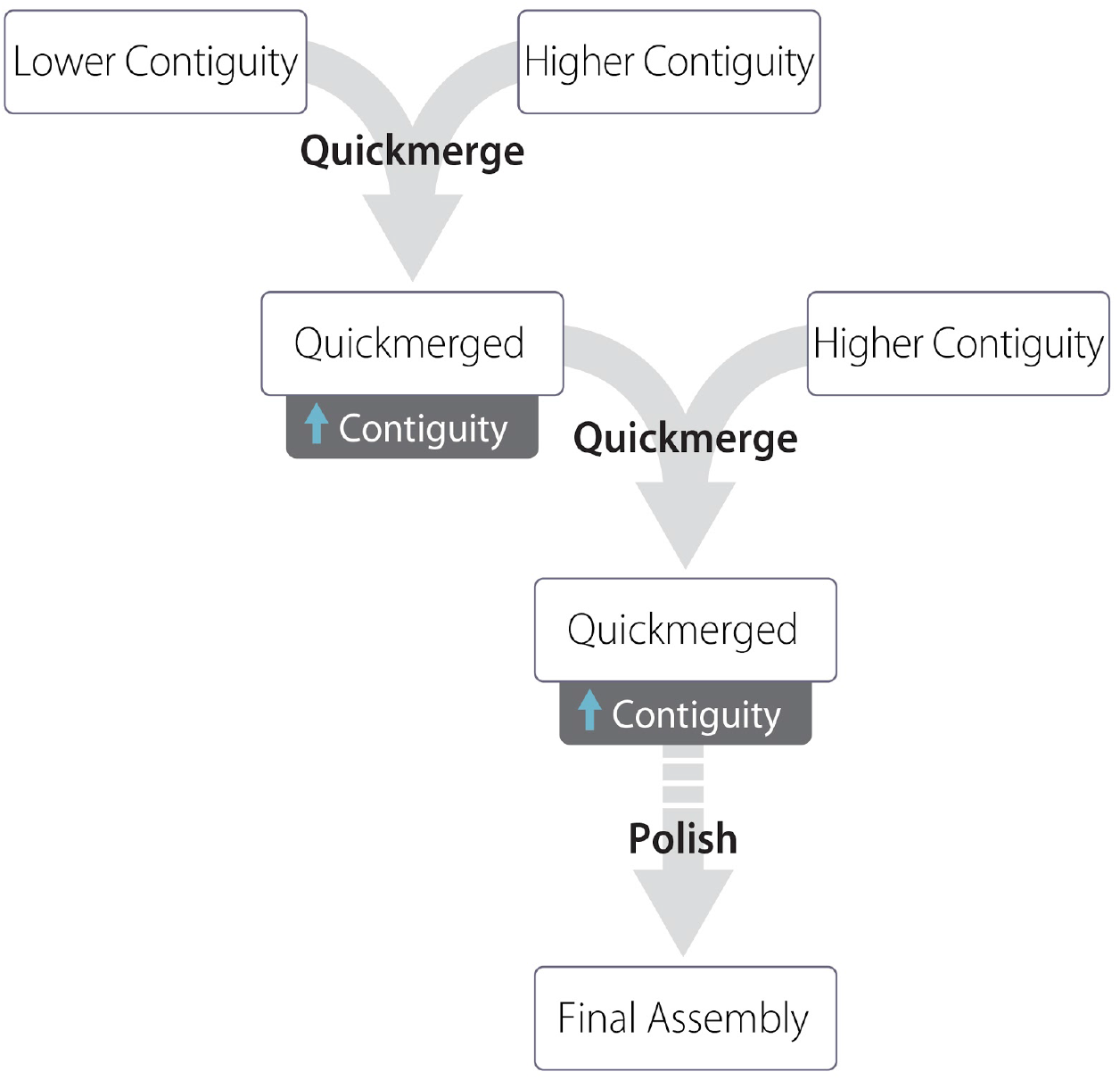
Assembly strategy used in this manuscript. A lower-contiguity assembly (Canu) is merged with a higher-contiguity assembly (DBG2OLC). The resulting assembly is again merged with the DBG2OLC assembly. The genome is then polished one or more times, here with nanopolish followed by Pilon.

### Assembly polishing

Assembly polishing was performed two ways. First, *nanopolish* version 0.7.1 (Loman, Quick, and Simpson 2015) was run using the recommended settings with reads longer than 1 kb. Prior to running nanopolish, the merged genome assembly was indexed using bwa, and ONT reads were aligned to the genome using bwa mem. The resulting bam file was then sorted, indexed, and filtered for alignments of size larger than 1 kb using samtools 1.3. Nanopolish was then run with a minimum candidate frequency of 0.1. Following nanopolish, we polished the assembly twice with Pilon (Walker et al. 2014) v1.16. For Pilon, we aligned 44x 150-bp and 67x 100-bp Illumina paired-end reads to the assembly using bowtie2 (Langmead and Salzberg 2012) and then ran Pilon on the sorted bam files using default settings.

### Bionano scaffolding

Bionano optical map data was collected following Chakraborty *et al*. (Chakraborty et al. 2018). ISO1 embryos less than 12 h old were collected on apple juice/agar Petri dishes, dechorionated using 50% bleach, rinsed with water, then stored at−80°C. DNA was extracted from frozen embryos using the Animal Tissue DNA Isolation kit (Bionano Genomics, San Diego, CA). Bionano Irys optical data was generated and assembled with IrysSolve 2.1 at Bionano Genomics. We then merged the Bionano assembly with the final merged assembly contigs using IrysSolve, retaining Bionano assembly features when the two assemblies disagreed.

### BUSCO analysis

We used BUSCO v1.22 to evaluate completeness and accuracy of all ISO1 assemblies (Simão et al. 2015). We used the Diptera database, which contains 2,799 highly conserved genes, to estimate assembly completeness.

### Assembly comparison and quality metrics

Assemblies were compared using alignment dot plots. For dot plots, assembled genomes were aligned to the reference *D. melanogaster* genome (r6.16) using nucmer (using options: minmatch 100, mincluster 1000, diagfactor 10, banded, diagdiff 5) (Kurtz et al. 2004). The resulting delta alignment files were used to create the dot plots either with mummerplot (using options: –fat –filter –ps) or ggplot. QUAST v4.5.0 was used to compare each generated assembly to the contig reference genome assembly *(D. melanogaster* r6.16) for completeness and errors. QUAST was run in gage mode on contigs larger than 1 kb and with the reference assembly as fragmented, as it was originally the scaffolded assembly (Salzberg et al. 2012; Gurevich et al. 2013). GAGE was run on two types of reference assembly: the full reference assembly, and only the part of the reference assembly comprising ordered and oriented contigs on chromosome arms (i.e., Muller elements) and the mitochondrial sequence. Quality score analysis was performed by aligning our Illumina short reads against each genome assembly using Bowtie. The number of variants were summed and divided by the total number of bases (following Berlin et al. 2015) and also by the total number of bases aligned: Perror=VariantsTotal Bases, where “Total bases” represents the total number of bases in the assembly or the total aligned bases (following Koren et al. 2017). These values are proxies for the probability of error and are used to calculate the QV score for each assembly according to the relation: QV=-10*log10Perror.

A modified pipeline originally published in McCoy et al. 2014 and modified following Berlin et al. 2015 and Koren et al. 2017 was used (this pipeline relies on release 5.57 naming conventions) for calculating total genome reconstruction. Nucmer was used to align each assembly to the FlyBase 5.57 reference, and then separated and merged by euchromatic and heterochromatic regions for each chromosome arm using BEDtools (Quinlan et al. 2010). Gene and TE reconstruction was completed similar to Berlin et al. (2015). For measuring the accuracy of gene models and TE reconstruction, both releases 5.57 and 6.16 of the reference were used. Fasta files containing all genes and separately, TEs for both releases were downloaded from FlyBase. In total, 17,730 gene models and 5,392 transposons from release 6.16, as well as 17,294 gene models and 5,409 transposons from release 5.57 were aligned to each genome assembly independently using nucmer. Nucmer was used to align each gene and transposon fasta file to each assembly independently. Alignments greater than 0%, 99% and equal to 100% similarity were reported.

### Structural variant detection

Large (>100 bp) SVs were detected by aligning the Bionano scaffolds to the FlyBase (dos Santos et al. 2015) reference assembly using MUMmer v3.23 (nucmer-maxmatch) (Kurtz et al. 2004) and then annotating the disagreements between the assemblies as indels and duplications using SVMU v0.2beta (commit 4e65e95) (Chakraborty et al. 2018). Insertions overlapping with RepeatMasker (Repeatmasker 4.0.7) annotated TEs were annotated as TE insertions. SVs were validated with at least two ONT reads spanning the entire genomic feature containing the SVs plus 200 bp on both sides of the SV. TEs were inferred to be segregating when corrected long reads supporting the TE insertions were contradicted by other reads showing absence of the TE. For validation with spanning long reads, we aligned Canu-corrected ONT reads to the FlyBase and Bionano assemblies using BLASR (v 1.3.1) (M. J. Chaisson and Tesler 2012) and the sorted alignment bam files were visually examined with IGV (Thorvaldsdóttir, Robinson, and Mesirov 2013).

### Mitochondrial genome identification

The mitochondrial genome was identified by using BLAST (Altschul et al. 1990) to compare our final assembly (the Bionano assembly) against the mitochondrial genome from r6.16 of the *D. melanogaster* genome. A single contig was identified (tig00000438_pilon_pilon_obj) with 99% identity to the reference mitochondrial genome. This contig contained two copies of the mitochondrial genome in tandem (assembly duplications of circular genomes are not uncommon when doing assembly with long-read data), therefore the first 16,806 and last 2,104 nucleotides were removed from the contig. We used MITOS (Bernt et al. 2013) with default settings, metazoan reference, and invertebrate genetic code to annotate both the reference mitochondrial genome and our assembled mitochondrial genome.

### Data availability

The Illumina and basecalled ONT data generated in this study has been uploaded to the National Center for Biotechnology Information (https://www.ncbi.nlm.nih.gov/) under bioproject PRJNA433573. Genomes assembled in this study are available at https://github.com/danrdanny/Nanopore_ISO1. Releases 6.16 and 5.57 of the D. melanogaster genome used in this study are available on FlyBase (http://www.flybase.org). Bioinformatic scripts used in this pipeline are available at https://github.com/esolares/DMI1. Original data underlying this manuscript can be accessed from the Stowers Original Data Repository at http://www.stowers.org/research/publications/libpb-1268.

## RESULTS

### Sequencing results

A 1D sequencing library was loaded onto a release 9.5 flow cell and run for approximately 24 hours (see Methods), generating a total of 663,784 reads (Table 1). Base calling was performed with Albacore 2.0.2, with 593,354 (89%) of all reads marked as “pass” (reads having a quality score ≥7) and an average fragment length of 7,122 bp (Figure 2). The read N50 for those that passed filter was 11,840, with 41 reads longer than 50 kb and a maximum read length of 379,978 bp (Figure S1).

**Table 1.** Statistics of reads used for genome assembly, only reads with quality scores ≥ 7 were used. A genome size of 140 Mb was used for all calculations.

|  |  |
| --- | --- |
| Total Reads | 593,354 |
| Avg Read Length | 7,122 bp |
| Total Bases Sequenced | 4,184,159,334 |
| Genome Coverage | 30.2x |
| Reads > 1 kb | 530,466 |
| Genome Coverage in Reads > 1 kb | 29.9x |
| Reads > 10 kb | 145,634 |
| Genome Coverage in reads > 10 kb | 17.5x |

**Figure 2.**
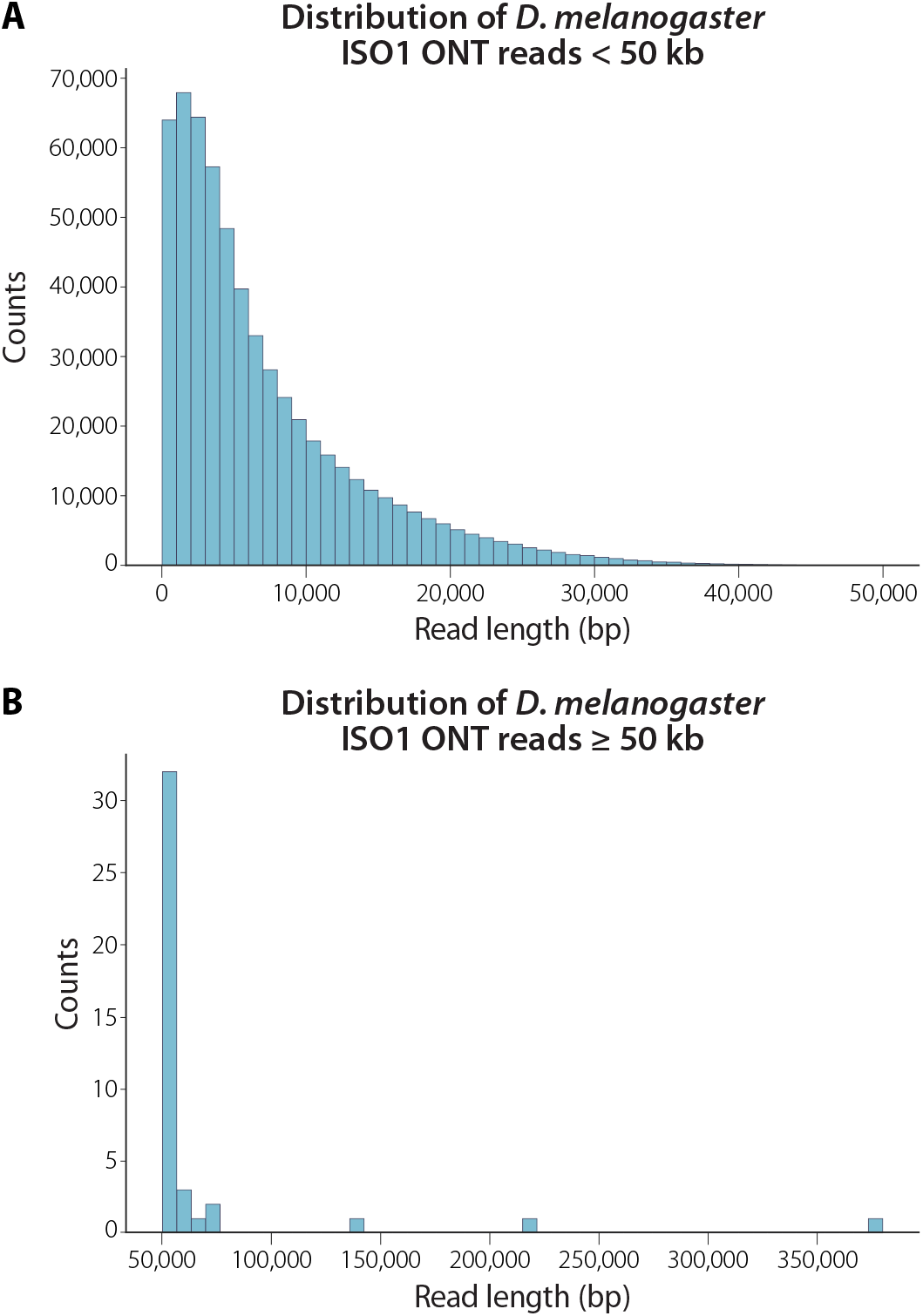
Read length distribution for reads with quality scores greater than or equal to 7. Length is the sequence length after base calling by Albacore, not the length that aligned to the genome. **(A)** Distribution of read lengths less than 50 kb. **(B)** Distribution of reads 50 kb or greater. The longest read that passed quality filtering was 380 kb.

### Genome assembly

#### ONT-only assembly using minimap/miniasm

To evaluate ONT data for *de novo* genome assembly, we performed an ONT-only assembly using minimap and miniasm (Li 2016). Together, these programs allow for rapid assembly and analysis of long, error-prone reads. We generated an assembly with a total size of 132 Mb, with 208 contigs and a contig N50 of 3.8 Mb (N50 is the length of the contig such that 50% of the genome is contained within contigs that are equal to or longer than this contig) (Table S2). Evaluation of an alignment dot plot between this assembly and the *D. melanogaster* reference genome revealed high correspondence between our assembly and the reference genome (Figure S2). However, BUSCO analysis of the minimap assembly found only 0.5% of expected single-copy genes present—much lower than the BUSCO score of 98.3% obtained from the current release of the *D. melanogaster* genome (Table 3). BUSCO analysis evaluates the presence of universal single-copy orthologs as a proxy of completeness. Such a low BUSCO score as found in the minimap assembly is unlikely to be measuring low completeness, but rather suggests a high rate of errors that disrupt the genes, making them difficult to properly assay.

#### ONT-only assemblies using Canu

We also generated an ONT-only assembly with Canu using only reads longer than 1 kb. Alignment dot plots for the assembled genome and the *D. melanogaster* reference genome also revealed large colinear blocks between this assembly and the reference, indicating only one large misassembly on chromosome 2L (this misassembly was broken prior to merging and polishing) (Figure 3A). The Canu assembly was marginally less contiguous (contig N50 = 3.0 Mb) than the minimap assembly, but resulted in a higher BUSCO score of 67.7% (Table S1). The errors in the Canu assembly and the low BUSCO score are consequences of inherently high error rate of ONT reads. However, due to the higher accuracy and completeness of the Canu assembly compared to the minimap assembly, we used the ONT-only Canu assembly for the remainder of our analysis.

**Figure 3.**
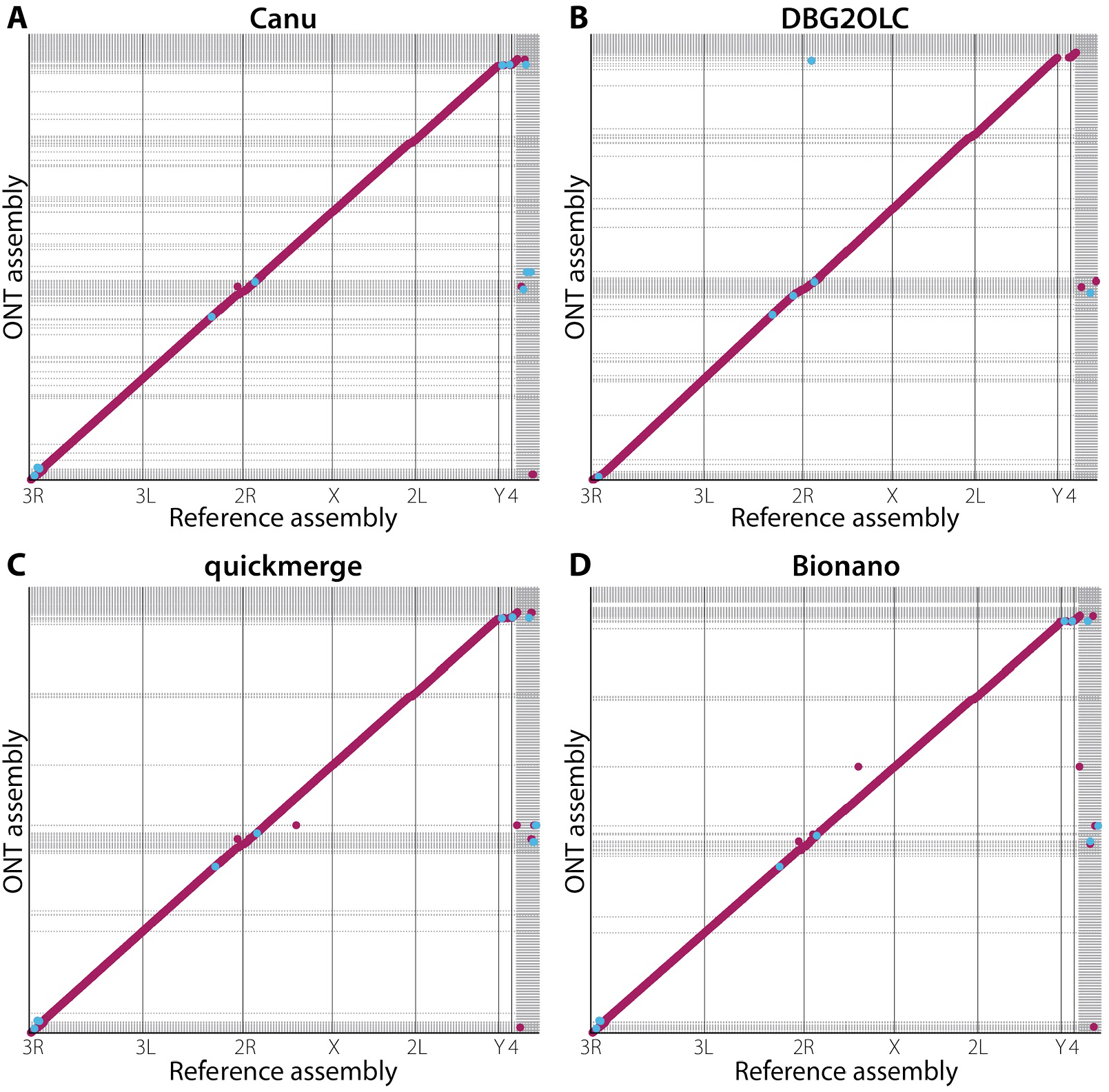
Dot plots showing colinearity of our assembled genomes with the current version of the *D. melanogaster* reference genome. Red dots represent regions where the assembly and the reference aligned in the same orientation; blue dots represent regions where the genomes are inverted with respect to one another. The vertical grid lines represent boundaries between chromosome scaffolds in the reference assembly. Horizontal grid lines represent boundaries between contig (A-C) or scaffolds (D) in the assemblies reported here. **(A)** Plot of the Canu-only assembly against the reference genome. **(B)** Plot of the hybrid DBG2OLC Nanopore and Illumina assembly against the reference. **(C)** Plot of merged DBG2OLC and Canu assemblies showing a more contiguous assembly than either of the component assemblies. **(D)** Bionano scaffolding of the merged assembly resolves additional gaps in the merged assembly.

#### Hybrid assembly using ONT and Illumina reads

Modest coverage assemblies of PacBio long reads can exhibit high contiguity when a hybrid assembly method involving Illumina reads is used (Chakraborty et al. 2016). Therefore, we examined whether such assembly contiguity improvements also occur when ONT long reads are supplemented with Illumina paired-end reads. We performed a hybrid assembly using DBG2OLC with the longest 30x ONT reads and contigs from a De Bruijn graph assembly constructed with 67x of Illumina paired-end data. To optimize the parameters, we performed a gridsearch on four parameters (AdaptiveTh, KmerSize, KCovTh and MinOverlap), yielding 36 genome assemblies. We used a range of values recommended by the authors for low-coverage assemblies and verified our KmerSize by looking at meryl’s kmer histogram and found it coincided with a value that represented a 99% fraction of all k-mers (http://kmer.sourceforge.net/wiki/index.php/Getting_Started_with_Meryl). We selected the best genome based on colinearity and largest N50. The best hybrid assembly was substantially more contiguous (contig N50 = 9.9 Mb) than the ONT-only Canu assembly (contig N50 = 3.0 Mb), had large blocks of contiguity with the reference (Figure 3B), yet had lower BUSCO scores (47.7% compared to the 67.7% observed in the Canu assembly) (Table 3).

#### Merging of Canu and DBG2OLC assemblies

We have previously shown that merging assemblies constructed using only PacBio long reads with hybrid assemblies constructed with PacBio long reads and Illumina paired-end short reads results in a considerably more contiguous assembly than either of the two component assemblies alone (Chakraborty et al. 2016). To examine the effect of assembly merging on assemblies created with ONT long reads, we merged the ONT-only Canu assembly with the DBG2OLC assembly with two rounds of quickmerge (see Methods). The merged assembly (contig N50 =18.6 Mb) was more contiguous than both the Canu and the DBG2OLC assemblies alone (Table 2), exhibiting large-scale colinearity with the reference genome as seen in the dot plot (Figure 3C). As expected, the BUSCO score of the merged assembly (58.2%) fell between the two component assemblies (Table 3).

**Table 2.** Genome assembly statistics. *Values are for scaffolds, not contigs.

| Name | Genome size (bp) | Contigs | Largest contig (bp) | N50 (bp) | L50 |
| --- | --- | --- | --- | --- | --- |
| FlyBase r6.16* | 142,573,024 | 2,442 | 27,905,053 | 21,485,538 | 3 |
| MiniMap | 131,856,353 | 208 | 16,991,501 | 3,866,686 | 9 |
| DBG2OLC | 131,359,678 | 339 | 13,129,070 | 9,907,730 | 6 |
| Canu | 139,205,737 | 295 | 14,326,064 | 2,971,262 | 11 |
| QuickMerge 2x | 138,130,519 | 250 | 25,434,901 | 18,616,266 | 4 |
| QM2x Nanopolish | 139,303,903 | 250 | 25,367,201 | 18,818,677 | 4 |
| QM2x NP + Pilon x2 | 140,153,080 | 250 | 25,783,280 | 18,923,871 | 4 |
| QuickMerge 2x Bionano | 142,817,829 | 231 | 28,580,427 | 21,305,147 | 3 |

### Assembly quality

#### Polishing

A high number of SNP and indel polymorphisms in all of our assemblies is consistent with other *de novo* assemblies created with noisy long reads with high error rates (Loman, Quick, and Simpson 2015; Chakraborty et al. 2016). Such errors typically lead to error-ridden genic sequence that can give the appearance that many important genes are missing (Table 3; Table S1). Many of these errors can be fixed via assembly polishing, and a number of assembly polishing tools exist. We chose to employ two: nanopolish, which performs consensus-based error correction using ONT reads (Loman, Quick, and Simpson 2015), and pilon, which performs error correction using Illumina reads (Walker et al. 2014). This approach of applying long-read consensus correction followed by short-read polishing has resulted in BUSCO scores comparable to that of the FlyBase reference genome (e.g. Chakraborty et al. 2016 and Chakraborty et al. 2018).

**Table 3.** Busco scores demonstrating genome quality before or after polishing.

| Name | Single copy | Duplicate | Fragmented | Missing | Complete |
| --- | --- | --- | --- | --- | --- |
| FlyBase r6.16 | 2,749 (98.2%) | 14 (0.5%) | 22 (0.8%) | 14 (0.5%) | 2,763 |
| MiniMap | 14 (0.5%) | 0 (0.0%) | 31 (1.1%) | 2,754 (98.4%) | 14 |
| DBG2OLC | 1,332 (47.6%) | 3 (0.1%) | 557 (19.9%) | 907 (32.4%) | 1,335 |
| Canu | 1,884 (67.3%) | 11 (0.2%) | 557 (19.9%) | 347 (12.4%) | 1,895 |
| QuickMerge (QM) 2x | 1,623 (58.0%) | 6 (0.3%) | 560 (20.0%) | 610 (21.8%) | 1,629 |
| QM 2x Nanopolish (NP) | 2,189 (78.2%) | 8 (0.3%) | 400 (14.3%) | 202 (7.2%) | 2,197 |
| QM 2x NP + Pilon x2 | 2,726 (97.4%) | 14 (0.5%) | 39 (1.6%) | 20 (0.7%) | 2,740 |
| QM 2x Pilon x2 | 2,718 (97.1%) | 14 (0.5%) | 45 (1.4%) | 22 (0.8%) | 2,732 |
| QM 2x Bionano | 2,715 (97.0%) | 15 (0.5%) | 40 (1.4%) | 29 (1.0%) | 2,730 |
| QM 2x Bionano All | 2,720 (97.2%) | 16 (0.6%) | 40 (1.4%) | 23 (0.8%) | 2,736 |

To evaluate this approach for ONT data, we evaluated three different polishing approaches: one using nanopolish alone, one using only two rounds of pilon, and one using one round of nanopolish followed by two rounds of pilon (see Methods). Applying nanopolish alone recovered only 79%, 79.2%, and 78.5% complete BUSCOs for the hybrid, ONT-only, and the merged assemblies, respectively (Table S2), suggesting that polishing with ONT reads alone only partially improved assembly quality. On the other hand, polishing all three assemblies twice with pilon alone fixed a large number of errors as evidenced by improved BUSCO scores of the resulting assemblies (Simão et al. 2015) (Table 3). The merged assembly was polished once with nanopolish, then twice with pilon, resulting in nearly all complete BUSCOs (97.9%) being present in our polished assembly, comparable to that of the reference assembly (98.3%) (Table S2). Variation in BUSCO scores generally tracked the method of polishing more than the type of assembly. However, the hybrid assembly did recover a slightly different subset of BUSCOs than the ONT-only assembly, leading to a slightly higher number of BUSCOs being recovered in the final merged assembly.

#### QUAST/GAGE metrics of quality

The QUAST output comparing the Canu, DBG2OLC, merged, and Bionano genome assemblies against the *D. melanogaster* reference shows that the quickmerge assembly resulted in intermediate error rates and discordance as compared to the component assemblies (Figure 4). All four assemblies exhibited approximately the same number of SNP and indel errors less than 5 bp, whereas the DBG2OLC assembly resulted in approximately 20% more indels greater than 5 bp in size than the Canu assembly (Figure 4H-I, Table S3, Table S4). Among ONT assemblies, the Canu assembly exhibited superior accuracy and fewer misassemblies, although our quickmerge assembly was nearly as good. When mismatches are measured on a phred scale, the ONT assemblies range in quality from 32 to 33, which is lower than the score of 37 from a PacBio assembly with threefold more data (Koren et al. 2017) (Table S3, Table S4). Canu also exhibited the lowest raw contiguity for an ONT assembly as measured by N50/NG50 and L50/LG50. The N50 for our merged assembly (19 Mb) exceeded that of Canu (3 Mb) and was nearly double that of DBG2OLC (10 Mb), while maintaining large-scale colinearity with the reference (Figure 3C). However, Canu outperforms the merging approach when considering the contiguity of error-free alignment blocks as demonstrated by the NA50 and NGA50 for Canu, which are greater than that of quickmerge (Table S3). We also observe that half of the assemblies accrue in fewer error-free alignments for Canu than for quickmerge, resulting in slightly lower values in LA50 and LGA50 for Canu than quickmerge (Table S3). As a consequence, when evaluating which approach to employ, we advise users to carefully weigh the tradeoffs between large-scale contiguity and local misassembly and sequence errors and to tailor their decision to the biological questions being addressed. For example, in creating a reference for QTL mapping, there might be a strong preference for high long-range contiguity even at the cost of local misassemblies. On the other hand, when avoiding misassemblies is paramount (as might be the case when characterizing the structure of individual loci), one could make the argument in favor of less long-range contiguity in exchange for fewer misassemblies.

**Figure 4.**
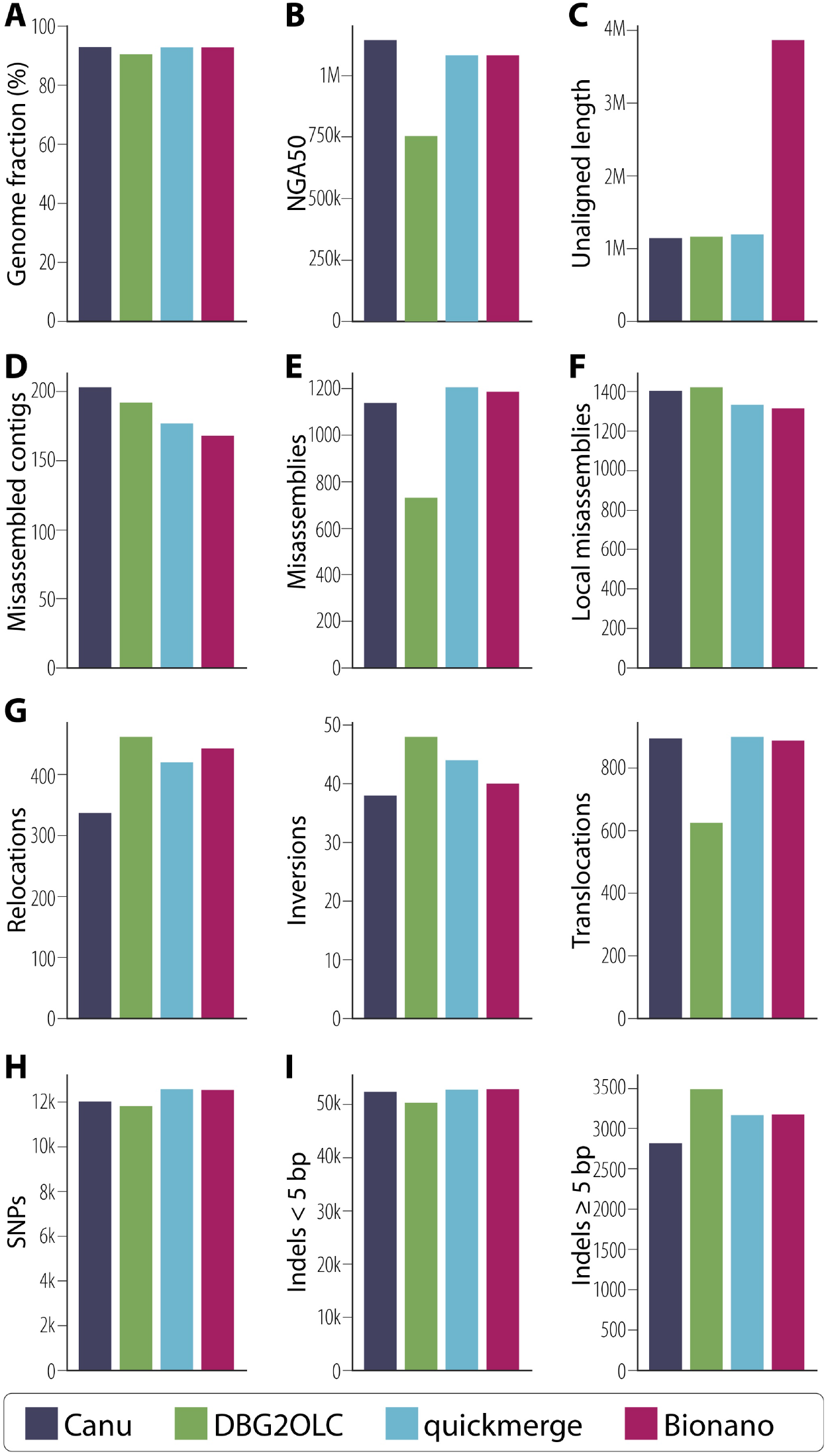
QUAST was used to compare each assembly to the *D. melanogaster* reference genome with selected statistics presented here. **(A)** Greater than 90% of bases in the reference genome were aligned to each of our four assemblies. **(B)** The contiguity of assembly blocks aligned to the reference. **(C)** Total unaligned length includes contigs that did not align to the reference as well as unaligned sequence of partially aligned contigs. **(D)** The number of contigs that contain misassemblies in which flanking sequences are 1 kb apart, flanking sequences overlap by 1 kb or more, or flanking sequences align to different reference scaffolds. **(E)** Total count of misassemblies as described in (D). **(F)** Local misassemblies includes those positions in which a gap or overlap between flanking sequence is less than 1 kb [(D) and (E) show those greater than 1 kb] and larger than the maximum indel length of 85 bp on the same reference genome scaffold. **(G)** Misassemblies can be subdivided into relocations (a single assembled contig aligns to the same reference scaffold but in pieces at least 1 kb apart), inversions (at least one part of a single assembled contig aligns to the reference in an inverted orientation), or translocations (at least one part of a single assembled contig aligns to two different reference scaffolds). Not all misassemblies are captured in these three categories. **(H)** Total SNPs per assembly are shown and were not significantly different among assemblies. **(I)** Indels per 100 kb can be divided into small indels (those <5 bp) and large indels (≥ 5 bp). Indels >85 bp are considered misassemblies and are shown in panels D, E, or F.

Finally, the low-coverage ONT dataset presented here performed surprisingly well compared to a PacBio dataset three times larger (Kim et al. 2018; Koren et al. 2017) in terms of contiguity (N50, NA50, L50, LA50), completeness (genome fraction), and accuracy (identity, SNPs, InDels, translocations, etc.) (Table S3). In all measures, the merged assembly performed almost as well as the much higher coverage PacBio assembly from Koren et al. 2017. Importantly, however, the GAGE corrected N50 of Koren et al. 2017 was substantially larger than the ONT assemblies, likely due to its superior coverage (Table S3, Table S4).

#### Completeness: reconstructing genes, TEs, and chromosome arms

We next evaluated the completeness of the assemblies by assessing the reconstruction of elements in the genome, including genes, TEs, and chromosome arms. For all following metrics, Canu and the final Quickmerge assembly performed similar to one another, with DBG2OLC performing somewhat worse. As a result, remaining comparisons will be made between the final Quickmerge ONT assembly and the PacBio assembly produced by Koren et al. 2017, which we use as a standard of comparison.

Our final assembly contained 96.37% of release 6.16 genes greater than 99% complete as compared to 98.4% for the Koren et al. 2017 assembly (Table S5). However, our final assembly was able to reconstruct only 72.71% of gene models compared to 88.14% for the Koren et al. assembly when only 100% complete gene models are considered (Table S5). Transposon reconstruction followed the same pattern with a 6.66% and a 15.51% difference in counts between the two assemblies for those with greater than 99% identity and complete reconstruction, respectively, with both favoring the higher-coverage PacBio assembly.

Muller element (chromosome arm) reconstruction was more similar between the ONT and PacBio assemblies for the euchromatic regions, with an average difference of 0.47% and a maximum difference of 0.94% in the X chromosome (Table S5). The largest difference was in the number of alignments. Bionano and quickmerge assemblies covered the Muller elements in fewer segments than the Koren et al. 2015 assembly, though they exhibited less total chromosome coverage. The differences were more apparent in the heterochromatic scaffolds, as our final assembly contained less coverage than the Koren et al. 2017 assembly, with an average deficit of 4.44% coverage and a maximum deficit of 7.89% across all heterochromatic scaffolds linked to Muller elements (Table S6). The difference in the Y chromosome was the largest, as expected, for our relatively low coverage of mixed sex flies compared to the high coverage of all males for Koren et al. 2017.

#### Base quality

We aligned Illumina short reads to measure the sequencing error rate following Berlin et al. 2015. The proportion of reads successfully mapped to assemblies varied from 93.5% to 94.6%. The aligned reads resulted in QV scores ranging from 40.2 to 41.1. A slightly different approach used by Koren et al. 2017 yielded very similar QV scores in the range of 40.1-40.2 (Table S7), suggesting that our assembly was of relatively high quality in areas where reads align well.

### Bionano scaffolding

Bionano fragments, which are substantially longer than ONT reads, can be used for scaffolding contigs across repetitive regions. We generated 81,046 raw Bionano fragments (20.5 Gb) with an average fragment length of 253 kb. To use these fragments for scaffolding, we first created a Bionano assembly (509 maps, haploid assembly size = 145 Mb) using 78,397 noise-rescaled fragments (19.9 Gb, mean fragment length 253 kb). At 145 Mb, the Bionano assembly is comparable to the reference assembly size (144 Mb). The Bionano maps were used to scaffold the merged assembly. The resulting scaffolded assembly was more contiguous (N50 = 21.3 Mb) than the unscaffolded contigs without substantially changing the number of SNP and indel errors (Figure 3D; Tables S1, S3-S4). Bionano scaffolding of our assembled genome did not change BUSCO scores (Table 3).

### Structural variants

One advantage of high quality *de novo* assemblies is that they permit comprehensive detection of large (>100 bp) SVs. Highly contiguous assemblies, such as the one generated here, allow comparisons between two or more assemblies, revealing novel SVs and facilitating the study of their functional and evolutionary significance. Although we sequenced the reference genome stock, structural differences between our stock and the published assembly are expected due to error, but also to new mutations— especially for TEs, which are the most dynamic structural components of the genome. Indeed, such mutations have been observed in substrains of ISO1 before (Zakharenko, Kovalenko, and Mai 2007; Moschetti et al. 2010; Rahman et al. 2015). Similarly, our assembly revealed the presence of 34 new TE insertions and 12 TE losses compared to the reference genome assembly. Among the 34 TE insertions, 50% (17/34) are LTR TEs comprising chiefly of *copia* (5/17) and *roo* elements (6/17). Interestingly, 29% (10/34) of the TE insertions are defective *hobo* elements that lack an average of 1.7 kb of sequence (base pairs 905-2510 of full-length *hobo* are absent) from the middle segment of the element encoding the transposase. However, alignment of long reads to the assembly regions harboring the TE insertions revealed that 6/34 insertions are not fixed, but are segregating in the strain (Table S2). The high insertion rate of the LTR and *hobo* elements in this single strain mirrors the recent spread of LTR and *hobo* elements in *D. melanogaster* populations (Pascual and Periquet 1991; Periquet et al. 1994; Bowen and McDonald 2001). As expected, the majority (27/34) of the new TE insertions are located within introns, because insertions within exons generally result in gene disruption. Nonetheless, we found five genes (*Ance, Pka-C1, CG31826*, *CG43446*, and *Ilp6)* in which new TEs have inserted within exons (Table S2). We also found 12 TEs present in the reference assembly that are missing from our final scaffolded assembly, among which six are LTR TEs (five *297* elements and one *roo* element). The high rate of insertion and loss of LTR elements underscores their dynamic evolutionary history in the *D. melanogaster* genome. Because TEs can be locally unique, the presence or absence of such events does not pose a fundamental limitation to assembly when reads are long enough to span the TEs. Consequently, we predict most of these events to be new mutations rather than errors in assembly.

Additionally, we identified several duplications present in our assembly. For example, a tandem duplication of a ∼9-kb segment has created partial copies of the genes *infertile crescent (ifc)* and *little imaginal discs (lid)*. Another ∼2-kb tandem duplication has created partial copies of the genes *CG10137* and *CG33116* (Table S8). Apart from CNVs affecting single copy sequences, our assembly also uncovered copy number increases in tandem arrays with potential functional consequences. For example, we observed copy number increase in a tandem array of a 207-bp segment within the third exon of the chitin-binding protein gene *Muc26B*. While CNVs such as this have been challenging to identify and validate in the past, at least two Nanopore reads spanning this entire tandem array support the presence of a tandem duplication at this position (Figure 5). Unlike TEs, classifying such tandem events as errors stemming from shorter Sanger reads or an actual mutation is difficult without the access to the original material from which the Sanger data was derived.

**Figure 5.**
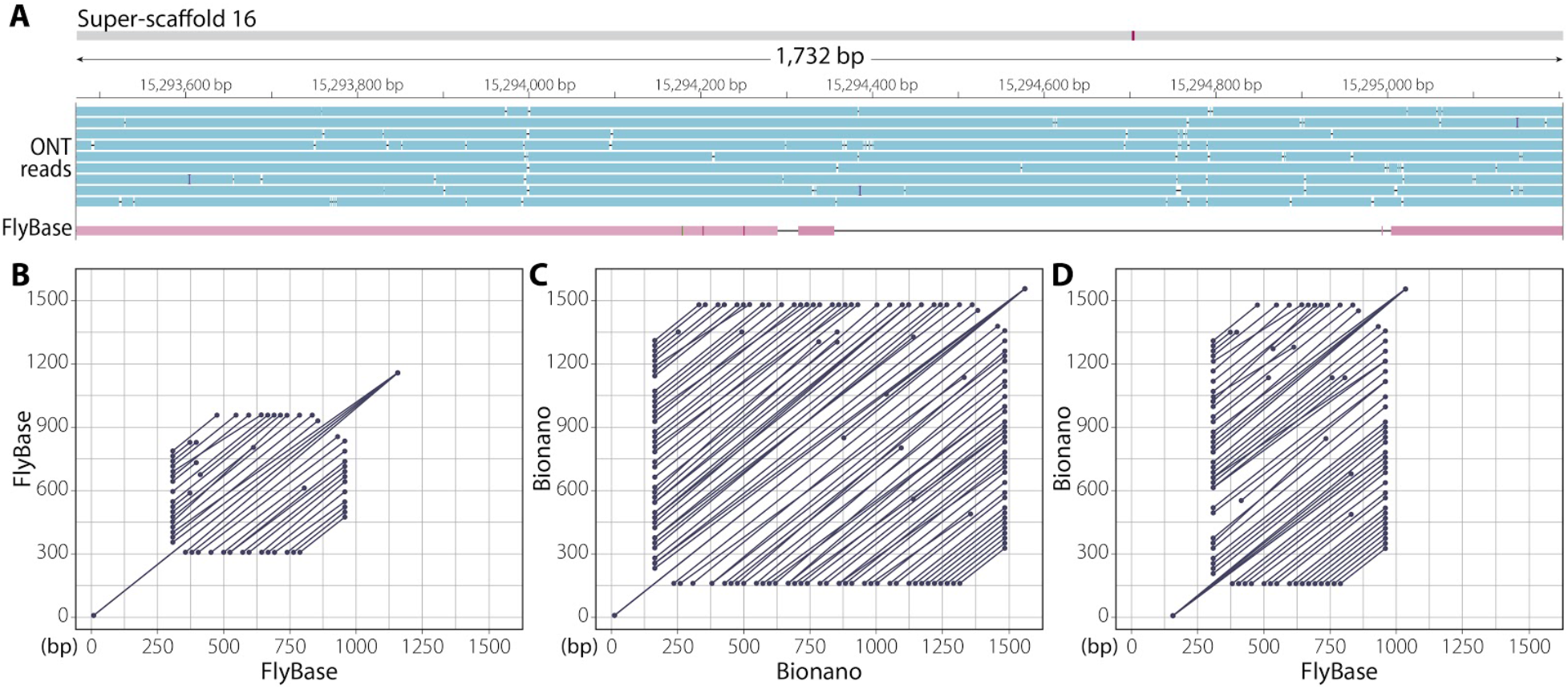
Copy number increase in a 207-bp tandem array located inside the third exon of *Muc26B*. **(A)** Three tracks showing Bionano assembly (top) with Nanopore long reads (blue) and reference (Flybase) assembly (red) aligned to it. The alignment gap in the reference assembly is due to the extra sequence copies in the Bionano assembly. **(B)** Alignment dot plot between the reference sequence possessing the tandem array to itself. **(C)** Alignment dot plot between the genomic region possessing the tandem array in the Bionano assembly to itself. As evidenced by the dot plot, the Bionano assembly has more repeats in this region than the reference assembly in panel B. **(D)** Alignment dot plot between the reference genomic region (x axis) shown in (B) and the corresponding Bionano genomic region (*y* axis) shown in (C).

### Mitochondrial genome identification and identity

After assembly, merging, polishing, and scaffolding, we used BLAST (Altschul et al. 1990) to identify the mitochondrial genome. Using the published *D. melanogaster* mitochondrial genome as the subject, we identified one 38,261-bp contig with nearly 100% identity to the reference mitochondrial genome. The first 16,806 and last 2,100 nucleotides of this contig were 99.6% identical to the reference mitochondrial genome, while the middle 19,228 nucleotides were 98% identical to the reference mitochondrial genome. This suggests that our assembled mitochondrial genome had been duplicated during assembly, which commonly occurs when assembling circular genomes using long sequencing reads. For the tandem assembled genomes, nearly all of the SNP and indel polymorphisms occurred in the last 4,000 nucleotides of the reference genome.

To determine if all genes and features that are present in the reference mitochondrial genome were present in our assembled genome, we annotated both genomes using MITOS (Bernt et al. 2013). Both assemblies were annotated nearly identically, with MITOS reporting that both assemblies were missing the origin of replication for the L region (OL) (Figure 6). The Nanopore assembly also resulted in two split genes not observed in the reference genome assembly: *nad4* and *nad6* were each annotated as two continuous genes rather than one single gene.

**Figure 6.**
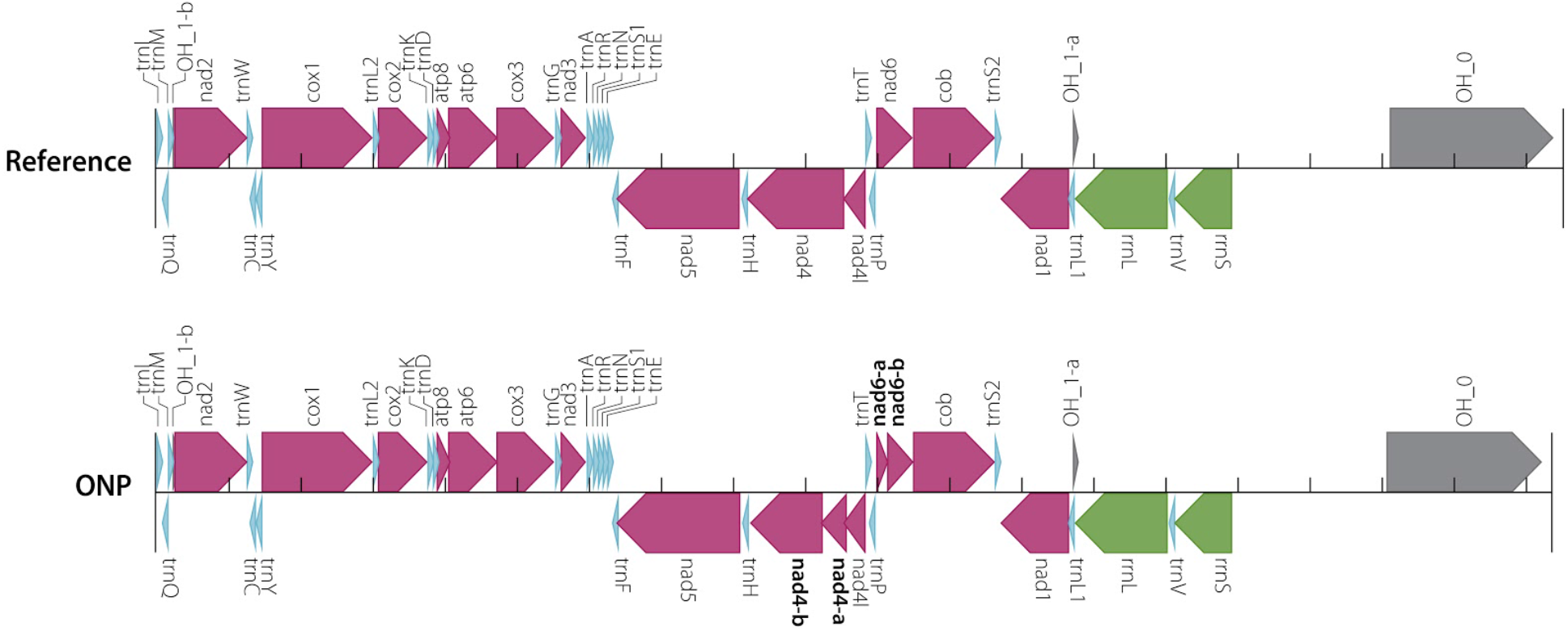
Mitochondrial genome annotations generated by MITOS. **(A)** Annotation of the reference mitochondrial genome. **(B)** Annotation of mitochondrial genome assembled in this project is identical to the reference except that *nad4* and *nad6* in the reference assembly were both annotated as two genes—*nad4* as *nad4-a* and *nad4-b*, and *nad6* as *nad6-a* and *nad6-b*.

## DISCUSSION

*Drosophila melanogaster* was the first genome assembled using a whole-genome shotgun (WGS) strategy (Myers et al. 2000; Adams et al. 2000). This successful proof-of-principle led to the prevalence of the WGS sequencing approach as a tool in virtually all subsequent metazoan genome assembly projects (Lander et al. 2001; Mouse Genome Sequencing Consortium et al. 2002; Goff et al. 2002; Yu et al. 2002; Aparicio et al. 2002; Gibbs et al. 2004; International Chicken Genome Sequencing Consortium 2004). While improvements in sequencing technology have led to a precipitous drop in the cost of sequencing (Anonymous 2018), stagnation and even regression of read lengths resulted in highly fragmented and incomplete assemblies (Alkan, Sajjadian, and Eichler 2011). Furthermore, complementary approaches (like hierarchical shotgun sequencing and other clone-based approaches) were required to obtain nearly complete and highly contiguous reference genomes, which adds complexity, cost, and time to assembly projects.

Short reads provided by next generation sequencing technologies present limitations to what a pure WGS assembly approach can accomplish (Alkan, Sajjadian, and Eichler 2011; Narzisi and Schatz 2015). The advent and development of long-read sequencing technologies has led to dramatic increases in read length, permitting assemblies that span previously recalcitrant repetitive regions. Early implementations of these technologies produced reads longer than previous short-read technologies yet still shorter than relatively common repeats, while the cost and error rate remained high compared to the short-read approaches. However, continued improvements in read length overcame many of these difficulties, permitting nearly complete, highly contiguous metazoan genome assemblies with only a WGS strategy (Kim et al. 2014; Berlin et al. 2015). Such approaches led to assemblies comparable in completeness and contiguity to release 6 of the *D. melanogaster* genome assembly (Hoskins et al. 2015) for approximately $10,000 USD (Chakraborty et al. 2016, 2018). However, even with the rapid development of these approaches, substantial capital investment in the form of expensive instrumentation and dedicated genome facility staff was required.

Here, we report an independent resequencing and assembly of the *D. melanogaster* reference strain, ISO1, for less than $1,000 USD in sequencing costs and without the need for extensive capital or personnel investment. The resulting assembly before scaffolding is comparable to release 6 of FlyBase in terms of contiguity and completeness (18.9 Mbp contig N50 and 97.1% complete single copy BUSCOs). We achieved this using 4.2 Gbp of sequence from a single Oxford Nanopore flow cell in conjunction with Illumina short-read data. Such reduction in complexity and cost permits a small team of scientists to shepherd a sequencing project from sample to near-reference-quality assembly in a relatively short amount of time. The addition of optical mapping data permitted ordering and orientation of the contigs, yielding an assembly nearly as contiguous as the published reference (scaffold N50 of 21.3 Mbp).

Comparing this assembly to the FlyBase reference genome shows that it is both accurate (21 mismatches/100 kbp, 36 indels/100 kbp) and colinear (Figure 3D, Figure 4, Table S3). Most of the small-scale differences are expected to be errors introduced by the noisy Oxford Nanopore reads that escaped correction via polishing. It is possible that some of these errors are SVs, which are expected to accumulate because the ISO1 stock has been maintained in the laboratory for approximately 350 generations (assuming 20 gen/year) since initial sequencing in 2000. This allows for the accumulation of new mutations by genetic drift, including ones reducing fitness (Assaf et al. 2017). Due to the high contiguity in the euchromatic region, our assembly facilitates detection of such euchromatic SVs. Several apparent “assembly errors” in our Bionano assembly are due to TE indels that are supported by spanning long reads. We found 28 homozygous euchromatic TE insertions, which are predominantly LTR and defective *hobo* elements, suggesting a high rate of euchromatic TE insertions (∼0.08 insertion/gen). That we observed a predominance of LTR and *hobo* elements among the new TE insertions mirrors their recent spread in *D. melanogaster* populations (Pascual and Periquet 1991; Periquet et al. 1994; Bowen and McDonald 2001; De Freitas Ortiz and Silva Loreto 2008). The abundance of defective *hobo* elements among the new insertions is particularly interesting given that these *hobo* elements lack the transposase enzyme necessary for mobilization.

Although most novel TE insertions were found in introns, five were found within exons: the 5’ UTR of *Ance, CG31826*, and *Ilp6*; the 3’ UTR of *Pka-C1*; and the coding region of *CG43446*. Similarly, TE loss primarily involved LTR TEs, including loss of TEs from the 5’ UTR of the genes *Snoo* and *CG1358*. We also observed copy number increases both in unique sequences as well as in tandem arrays (Table S8), with one duplication creating a new copy of the entire coding sequence of the gene *lid*. Collectively, our assembly provides a snapshot of ongoing genome structure evolution in a metazoan genome, which is often assumed to be approximately invariant for experimental genetics.

A crucial feature of this work is that it is performed in a strain used to generate one of the highest quality reference genomes available, ensuring that our inferences can be judged against a high-quality standard. This approach allowed us to demonstrate that assembly with modest amounts of long-molecule data paired with inexpensive short-read data can yield highly accurate and contiguous reference genomes with minimal expenditure of resources. This demonstration opens myriad opportunities for high-quality genomics in systems with limited resources for genome projects. Moreover, we can now conceive of studying entire populations with high-quality assemblies capable of resolving repetitive structural variants, something previously unattainable with short-read sequencing alone.

---

**Figure S1.**
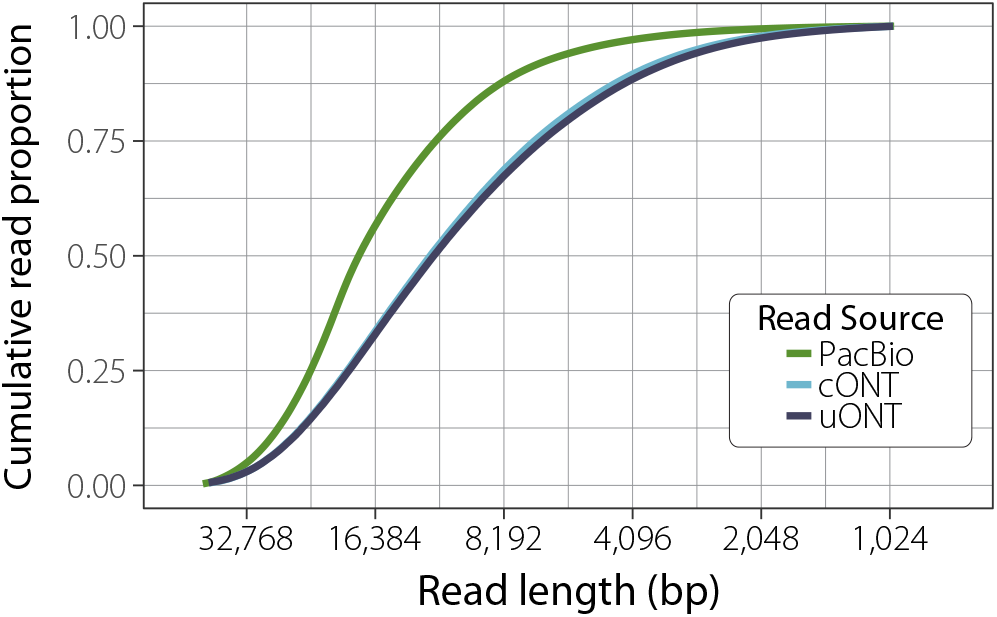
Proportion of corrected (cONT) and uncorrected (uONT) reads >1 kb used in this study compared to PacBio data collected for Chakraborty et al. (2018).

**Figure S2.**
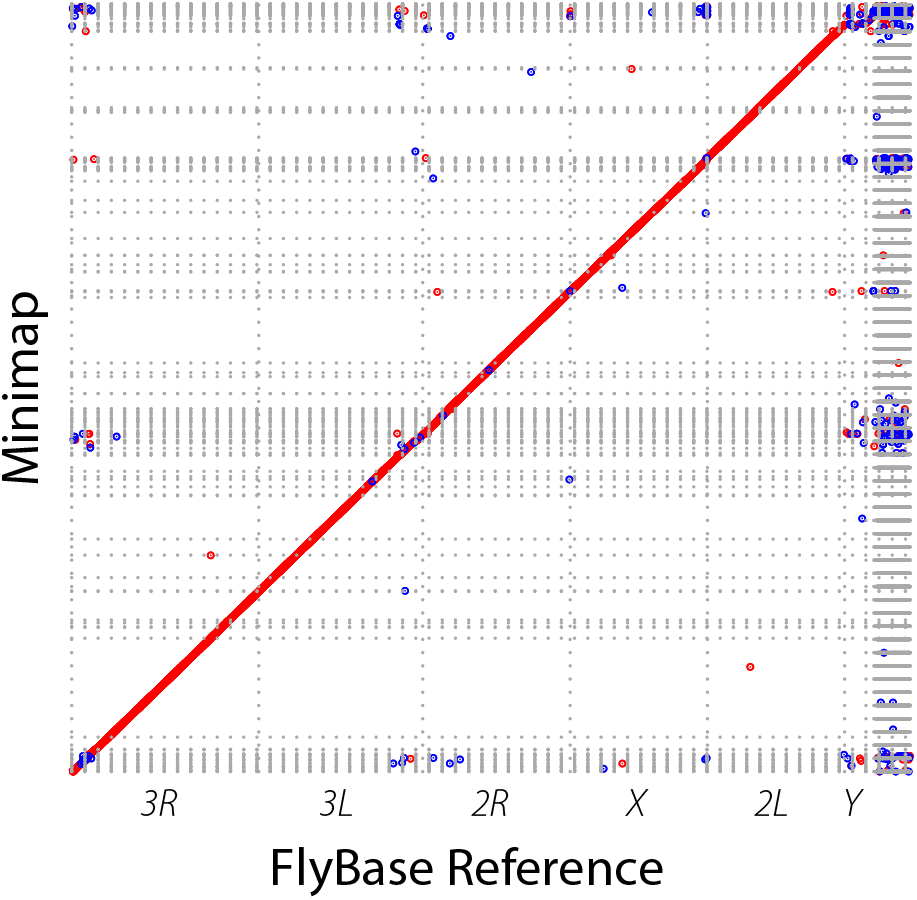
Dot plot of minimap assembly (*y* axis) compared to *D. melanogaster* reference assembly (*x* axis).

**Table S1.** BUSCO analyses for unpolished assemblies.

**Table S2.** BUSCO analyses for polished assemblies.

**Table S3.** GAGE statistics from QUAST for all release 6 contigs.

**Table S4.** GAGE statistics from QUAST for only Muller elements and the mitochondrial sequence from ISO1 release 6 contigs.

**Table S5.** Gene model and transposon completeness assessment following Berlin *et al*. (2015).

**Table S6.** Assembly completeness assessment for both heterochromatin and euchromatin using genome assembly alignments to reference 5.57, following McCoy *et al*. (2014) but modified by Berlin *et al*. (2015) and Koren *et al*. (2017).

**Table S7.** Quality analysis using QV scores following Koren *et al*. (2017) and Berlin *et al*. (2015).

**Table S8.** Detailed position information for SVs identified in this report.

## ACKNOWLEDGEMENTS

We would like to thank Christina Lin for assistance with the Bionano scaffolding, Melanie Oakes for help generating the Bionano data, and Angela Miller for assistance with editing and figure preparation. We thank Casey Bergman for providing examples of novel TEs in substrains of ISO1. DEM is part of the Oxford Nanopore Educational Initiative and has received free flow cells as part of that program. This material is based upon work supported by the National Science Foundation Bridge to the Doctorate Fellowship under Grant No. 1500284 and the National Science Foundation Graduate Research Fellowship under Grant No. DGE-1321846. MC is supported by US National Institutes of Health (NIH) grant R01 RR024862-01A6. JJE is supported by NIH grant R01GM123303-1 and University of California, Irvine setup funds. This work was made possible, in part, through access to the Genomics High-Throughput Facility Shared Resource of the Cancer Center Support Grant CA-62203 at the University of California, Irvine, and NIH shared-instrumentation grants 1S10RR025496-01, 1S10OD010794-01, and 1S10OD021718-01. RSH is an American Cancer Society Research Professor.

